# Inactivation of mitochondrial Complex I stimulates chloroplast ATPase in Physcomitrella (*Physcomitrium patens*)

**DOI:** 10.1101/2020.11.20.390153

**Authors:** Marco Mellon, Mattia Storti, Antoni Mateu Vera Vives, David M. Kramer, Alessandro Alboresi, Tomas Morosinotto

**Affiliations:** Department of Biology, University of Padova, 35121 Padova, Italy; MSU-DOE Plant Research Laboratory, Michigan State University, East Lansing, MI 48824, United States of America; Biochemistry and Molecular Biology, Michigan State University, East Lansing, MI 48824, United States of America

## Abstract

While light is the ultimate source of energy for photosynthetic organisms, mitochondrial respiration is still fundamental for supporting metabolism demand during the night or in heterotrophic tissues. Respiration is also important for the metabolism of photosynthetically active cells, acting as a sink for excess reduced molecules and source of substrates for anabolic pathways. In this work, we isolated Physcomitrella (*Physcomitrium patens*) plants with altered respiration by inactivating Complex I of the mitochondrial electron transport chain by independent targeting of two essential subunits. Results show that the inactivation of Complex I causes a strong growth impairment even in fully autotrophic conditions in tissues where all cells are photosynthetically active. Complex I mutants show major alterations in the stoichiometry of respiratory complexes while the composition of photosynthetic apparatus was substantially unaffected. Complex I mutants showed altered photosynthesis with higher yields of both Photosystems I and II. These are the consequence of a higher chloroplast ATPase activity that also caused a smaller ΔpH formation across thylakoid membranes as well as decreased photosynthetic control on cytochrome *b6f*, possibly to compensate for a deficit in ATP supply relative to demand in Complex I mutants. These results demonstrate that alteration of respiratory activity directly impacts photosynthesis in *P. patens* and that metabolic interaction between organelles is essential in their ability to use light energy for growth.

## Introduction

In photosynthetic organisms, sunlight powers the linear electron flow (LEF) from water to NADP^+^ catalysed by two photosystems (PS), PSII and PSI, cytochrome *b_6_f* and ATPase, ultimately generating NADPH and ATP to sustain cellular metabolism. In photosynthetic organisms, mitochondrial respiration is also active with its specific electron transport pathway, or oxidative phosphorylation (OXPHOS). OXPHOS transfers electrons from the substrates NADH and succinate to molecular oxygen through the activity of enzymatic complexes localized in the inner mitochondrial membrane, Complexes I, II, III, IV. This electron transfer is coupled to the generation of electrochemical transmembrane gradient that drives the synthesis of ATP through ATP-synthase, also called complex V. The NADH dehydrogenase complex (Complex I, CI) is the main site for electron insertion into the mitochondrial electron transport chain (mETC) and it can provide up to 40% of the protons for mitochondrial ATP formation (Watt et al., 2010; Braun et al., 2014). In plants mETC electrons can also follow alternative routes, bypassing CI via alternative NADH dehydrogenases (Lecler et al., 2012) and complex III / IV via the alternative terminal oxidase (AOX) (Dinant et al., 2001). These alternative routes partly uncouple the electron transport and the electrochemical transmembrane gradient, thus reducing the energy yield of respiration with beneficial effects especially in stress conditions (Zalutskaya et al., 2015), reducing ROS production from mETC (Møller, 2001; Vanlerberghe et al., 2020).

Respiration in photosynthetic organisms is essential to support energy demand during the night, in non-photosynthetic tissues such as roots or in developmental stages where photosynthesis is not active (e.g. seed germination). An increasing set of evidence, however, underlines the importance of respiration also for sustaining optimal photosynthetic activity (Joliot and Joliot, 2008; Noguchi and Yoshida, 2008; Bailleul et al., 2015) with a strong functional link between chloroplasts and mitochondria bioenergetic metabolism (Cardol et al., 2003; Dutilleul et al., 2003; Schönfeld et al., 2004). In the diatom *Phaeodactylum tricornutum* this was clearly shown by demonstrating that metabolites exchange between chloroplast and mitochondria is essential for carbon fixation (Bailleul et al., 2015). In another example of such a functional link, excess reducing power produced via photosynthesis can be routed to mitochondrial respiration, preventing over-reduction and eventual reactive oxygen species (ROS) production in the plastid (Noguchi and Yoshida, 2008; Zhang et al., 2012). AOX activity has also been shown to influence photosynthetic metabolism response to stresses when it consumes excess reductant while decreasing mitochondrial ATP synthesis (Cheung et al., 2015) and maintaining redox balance of plastoquinone pool (Yoshida and Noguchi, 2011; Vanlerberghe et al., 2020). Consistently, AOX protein level was shown to be linked to differences in chloroplast energetic balance, being induced under strong irradiance, suggesting the presence of regulation signals originated from photosynthetic ETC activity (Dahal et al., 2016).

Imbalances between the relative rates of production and consumption of ATP/NADPH utilization can lead to the build-up of reactive intermediates of electron transfer processes, driving the formation of harmful ROS (Eberhard et al., 2008; Li et al., 2009). Photosynthetic organisms evolved multiple regulatory mechanisms to balance light-dependent processes and metabolic exploitation of photosynthesis products. Examples of such mechanisms are the dissipation of excess excitation energy as heat (non-photochemical quenching, NPQ) or the photosynthetic control to reduce electron transport capacity at the level of cytochrome *b_6_f* and prevent over-reduction. Both mechanisms are activated by a decrease of lumenal pH, that represents a major signal for regulation of photosynthesis (Eberhard et al., 2008; Li et al., 2009).

Consistently with the essential role of mitochondrial respiration in plants metabolism, Knock-Out (KO) mutants completely depleted in complex II (Leon et al., 2007), complex III (Colas Des Francs-Small and Small, 2014) and complex IV (Radin et al., 2015) are not viable and so far only knock-down plants have been isolated and studied. Mutants completely lacking mitochondrial CI activity have instead been described in *Arabidopsis* (Kühn et al., 2015; Fromm et al., 2016a) as well as in *Nicotiana tabacum* (Vidal et al., 2007) where they showed a severe growth phenotype and alterations in germination, fertilization and pollen development (Fromm et al., 2016a). European mistletoe *Viscum album* can live without CI, but it is an obligate semi-parasite living on branches of trees, and thus its energy metabolism is expected to be remodelled (Senkler et al., 2018).

Differently from plants, a large number of respiratory mutants have instead been isolated in the green alga *Chlamydomonas reinhardtii* where they generally show strong phenotypes under heterotrophic conditions but grow similarly to WT under photo-autotrophic conditions (Salinas et al., 2014; Larosa et al., 2018).

These differences suggest that the role of respiration on cells metabolism adapted during plants evolution, motivating for the investigation of species that diverged at different times during evolution. Non-vascular plants like the moss Physcomitrella (*Physcomitrium patens*) diverged from vascular plants ancestors early after land colonization and thus their study allows highlighting the first adaptation to the new environmental conditions. To assess how the biological role of respiration adapted during the evolution of photosynthetic organisms in this work, we generated *P. patens* plants depleted of active respiratory CI and investigated effects on photosynthetic activity. Results show that the absence of CI stimulates photosynthetic transport caused by more active chloroplast ATPase with alteration of pmf and ΔpH across the thylakoid membranes and impairment of photosynthetic control.

## Results

Complex I (CI) in eukaryotes is composed of over 40 subunits and, among them, 14 are highly conserved across kingdoms (Ligas et al., 2019). Plant CI includes 9 additional subunits that form a carbonic anhydrase domain (Braun et al., 2014; Fromm et al., 2016a; Subrahmanian et al., 2016). *NDUFA5* and *NDUFB10* genes were selected to generate *P. patens* mutants depleted in CI based on two criteria: i) they encoded for conserved proteins known to be essential for CI activity in *C. reinhardtii, A. thaliana* or *H. sapiens* (Barbieri et al., 2011; Rak and Rustin, 2014); ii) they were present in a single copy in *P. patens* nuclear genome, facilitating the generation of mutants and ensuring that any eventual phenotype would be readily assessable. NDUFA5 is localized in the hydrophilic region of the complex binding the substrate NADH (Figure 1A) and it is required for assembly and stability of the matrix arm of CI in human mitochondria (Rak and Rustin, 2014). NDUFB10 deletion in *C. reinhardtii* impairs the assembly of the distal part of CI membrane module (Figure 1A), responsible for the proton pumping activity coupled with the electron transfer (Barbieri et al., 2011).

**Figure 1.**
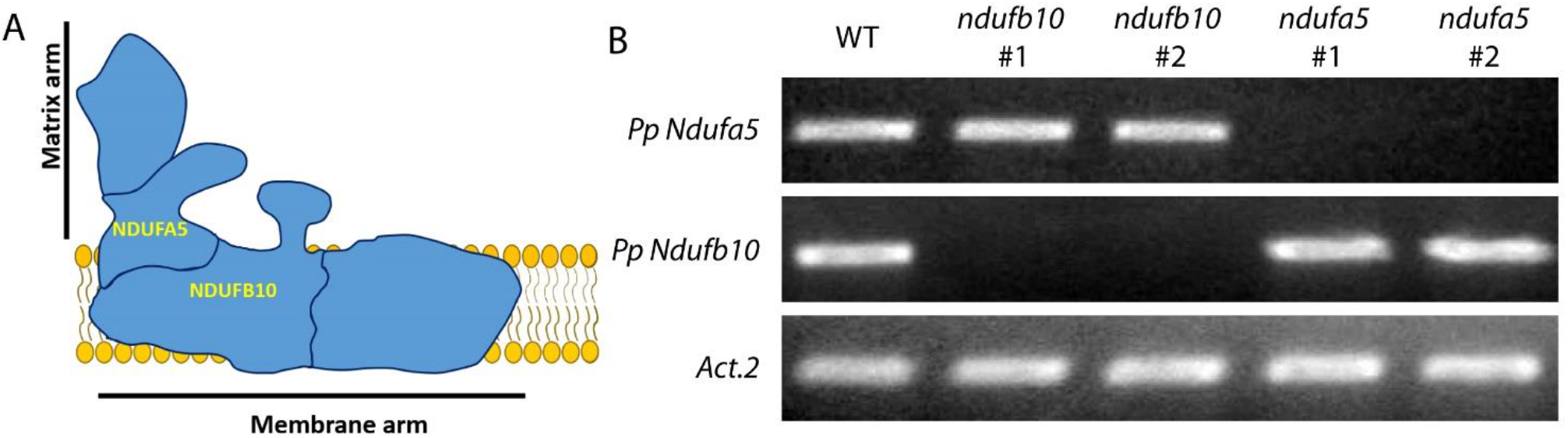
Generation of *P. patens* plants depleted in Complex I. A) Schematic representation of CI structure with NDUFA5 and NDUFB10 localization. B) RT-PCR to assess the accumulation of the mRNA encoding for *Ndufa5* and *Ndufb10* in two independent KO lines.

KO lines for both genes were generated by homologous recombination-mediated gene targeting (Figure S1A-B). Multiple independent KO lines for each gene were isolated and the insertion of DNA in the expected position of the genome was verified by PCR (Figure S1C). The loss of expression of the target gene was also confirmed by RT-PCR (Figure 1B). In the following results from one line per gene are reported, but at least 4 independent confirmed lines per gene were isolated per each genotype.

All confirmed KO plants showed strongly impaired growth (Figure 2) that was visible upon cultivation on a glucose enriched medium but also on a mineral media in fully autotrophic conditions. Remarkably the growth defect was observed also if plants were grown autotrophically under 24 hours of continuous illumination, thus avoiding any dark time when respiration is expected to be essential (Figure 2). Glucose presence in the media and continuous illumination stimulated a faster growth in WT plants while the mutants remained unaffected (Figure S2).

**Figure 2.**
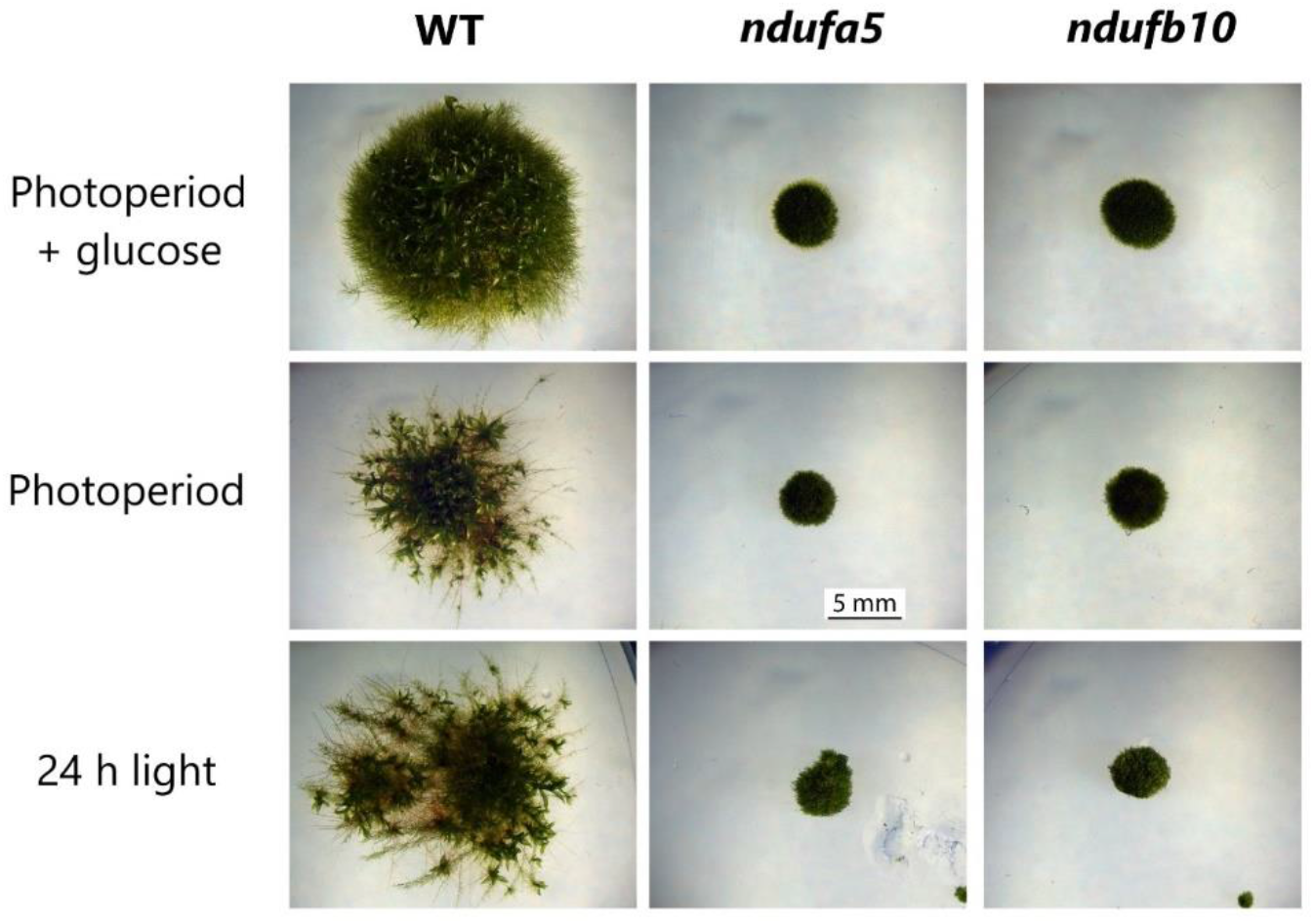
Impact of CI inactivation on *P. patens* growth. WT, *ndufb10* and *ndufa5* KO lines growth after 28 days under 16 d / 8 h dark photoperiod with/without the addition of 0.5% glucose in the media (top/middle respectively) or under 24 hours continuous illumination without glucose (bottom).

Impact of mutations on respiratory apparatus composition was assessed using specific antibodies (Figure 3A). NAD9, a CI core subunit localized in the Q module of the matrix arm, was completely missing in *ndufa5* mutant, consistently with recently reported evidence that NDUFA5 directly interacts with NAD9 during CI biogenesis (Ligas et al., 2019). NAD9 was instead present in *ndufb10* mosses suggesting that at least part of CI was still present, as observed for other analogous mutants (Ligas et al., 2019). Both KO plants showed significant alterations in other complexes of the respiratory apparatus. While CIV and CV contents were unaltered, CII and CIII were more abundant in the mutants as compared to WT plants by respectively ≈ 3 and 1.5 times. The most striking difference, however, was that AOX accumulated to approx. 10 times higher levels in *ndufa5* and *ndufb10* than in WT. All differences in protein accumulation were consistent between the two independent mutant lines.

**Figure 3.**
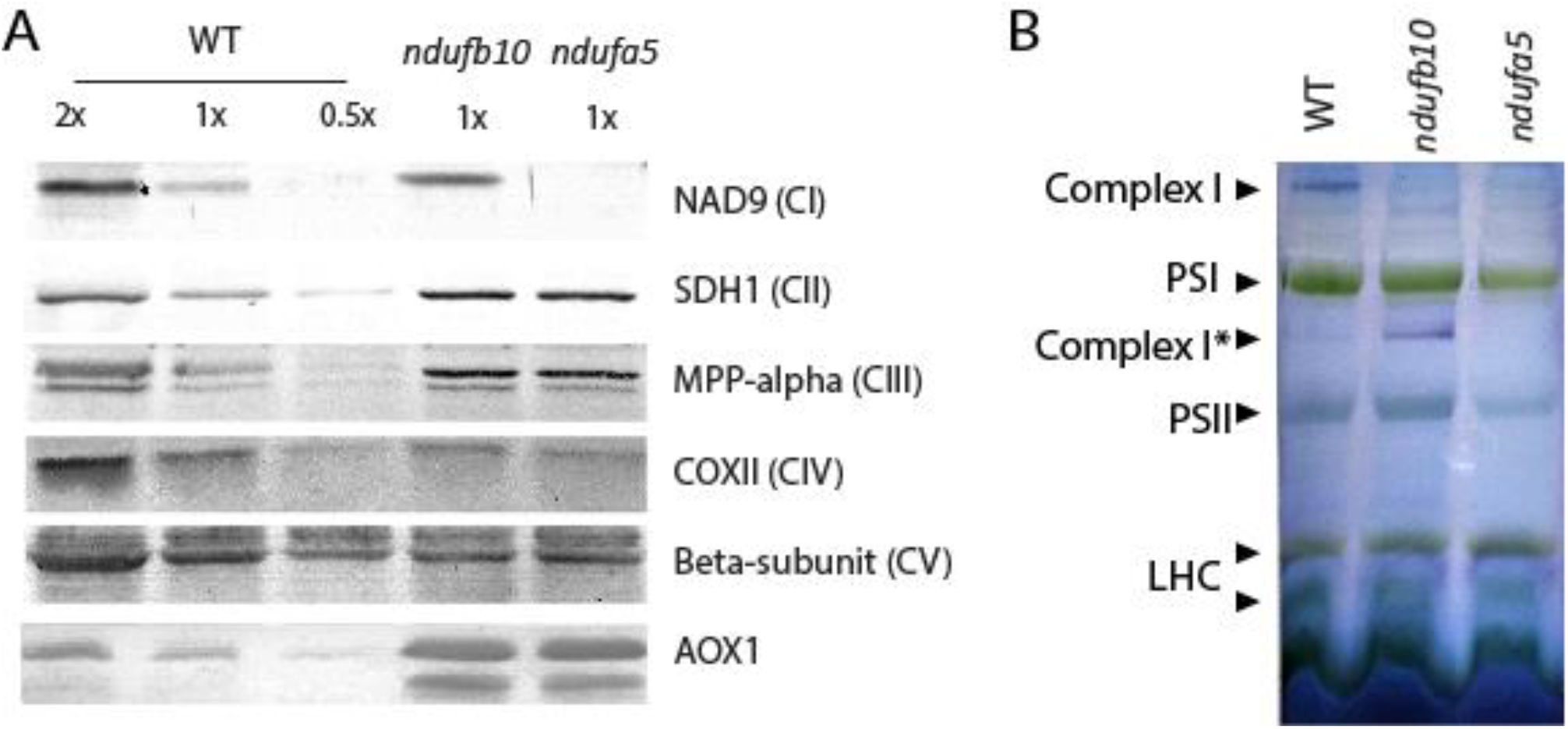
Impact of Complex I mutation on the respiratory apparatus composition and Complex I assembly. A) Immunoblot analysis using antibodies against various subunits of respiratory complexes. Total protein extracts were loaded, 1X samples correspond to 2 μg of total chlorophyll. B) BN-PAGE separation of crude organelles extracts. Following separation (see Figure S3), CI activity was detected with in-gel NADH/NBT staining. Membrane proteins from WT, *ndufb10* and *ndufa5* were solubilized with 1% of β-dodecyl maltoside (β-DM). Main complexes of thylakoid and mitochondria complex I are indicated on the profiles. Complex I* indicates an assembly intermediate of complex I present in *ndufb10*; PSI, Photosystem I; PSII, photosystem II; LHC, light-harvesting chlorophyll complexes.

The impact of mutations on CI was also assessed by blue native electrophoresis (BN-PAGE) on crude membrane extracts, containing both mitochondria and chloroplasts (Figure S3). The detection of CI enzymatic activity after electrophoresis showed that in both mutant lines the correct assembly of CI was compromised (Figure 3B). No CI activity was detectable in the case of *ndufa5*. In the *ndufb10* mutant, instead, the complete holoenzyme was absent but a partially assembled CI retaining the NADH dehydrogenase activity was detectable (Complex I*, Figure 3B) consistently with earlier results in other species (Barbieri et al., 2011).

The impact of mutations on respiratory activity was assessed measuring oxygen (O_2_) consumption rate. In both *ndufa5* and *ndufb10* plants, O_2_ consumption in the dark was higher than in WT (Figure 4). While this seems counterintuitive for mutants affected in a respiratory complex, this observation can be explained by the compensatory over-accumulation of CII, CIII and AOX. A similar increase in oxygen consumption was indeed observed also in *Arabidopsis* CI mutants (Kühn et al., 2015). This hypothesis is confirmed by the observation that O_2_ consumption activity in *ndufa5* and *ndufb10* was completely insensitive to the addition of rotenone, a specific CI inhibitor, that in WT plants instead reduced O_2_ consumption by ≈ 40 % (Figure 4A).

**Figure 4.**
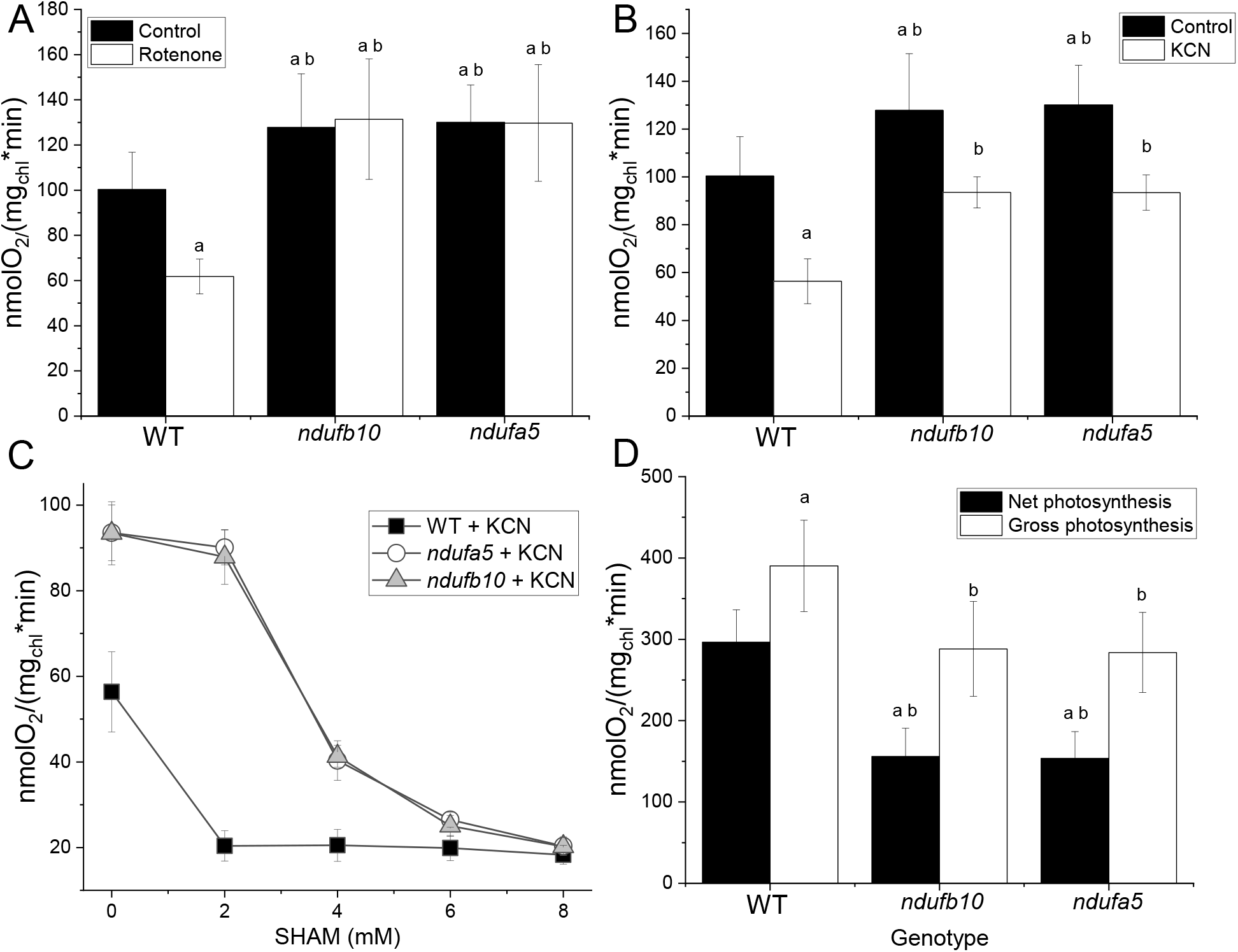
Oxygen consumption and evolution capacity in the *ndufb10* and *ndufa5*. A) Oxygen consumption was monitored with a Clark-type oxygen electrode on WT and *ndufb10* and *ndufa5* plants grown for 10 days in minimal medium. Mosses were maintained in the dark in the absence (black) or presence of 50 μM rotenone (white columns). B) Effect of cyanide (KCN) on O_2_ consumption. Controls without the inhibitor are in black, samples with addition 1mM KCN in white. In both panels “a” indicates a statistically significant difference compared to untreated WT, “b” significant difference compared to WT treated with the same inhibitor (one-way ANOVA, n > 5, p < 0.01). C) Effect of different concentration of SHAM on O_2_ consumption in the presence of 1mM KCN. WT is showed as black square, *ndufb10* as grey triangles, *ndufa5* as white circles. Average ± SD (n ≥ 5) are reported. D) Oxygen production of protonema illuminated with a light at 850 μmol photons m^-2^ s^-1^ measured with a Clark-type oxygen electrode. Net photosynthesis (black) is calculated directly from the O_2_ evolution rate during illumination. Gross photosynthesis (white) is calculated by adding net photosynthesis with oxygen consumption in the dark. In all panels average ± SD (n ≥ 5) is reported. “a” indicates statistically significant difference compared to WT net oxygen evolution, “b” indicates difference compared to WT gross O_2_ evolution (one-way ANOVA, n > 5, p < 0.01).

Oxygen consumption activity was similarly reduced by the addition of KCN, a Complex IV inhibitor, in both WT and mutants (Figure 4B). The significant residual activity still present was mostly attributable to the presence of alternative oxidases like AOX, as confirmed by the further decrease in O_2_ evolution induced by its specific inhibitor SHAM (Figure 4C). While 2 mM SHAM was a saturating dose for WT, a 4-fold higher dose was necessary to obtain a full inhibition in the mutants, consistent with their increased AOX content. Maximal photosynthetic O_2_ evolution activity was measured in the same samples upon exposition to saturating light. *ndufa5* and *ndufb10* KO mutants showed a reduction of ≈ 50 % (Figure 4D). At least a major fraction of this reduction can, however, be explained by the higher O_2_ consumption rate in the mutants than in WT.

### Impact of CI depletion on photosynthesis

Plants depleted of CI showed reduced photoautotrophic grow and alterations in O_2_ evolution activity, motivating a deeper investigation of the impact of the mutation on photosynthetic activity. Chlorophyll (Chl) content, Chl a/b and Chl/Car ratio was unchanged in mutant and WT plants (Table 1). Consistently, the accumulation of all main components of the photosynthetic apparatus (PSI, PSII, cyt *b6f*, ATPase) showed no major alterations in the mutants except for a slight increase in chloroplast ATPase content (Figure S4). Subunits involved in regulatory mechanisms of photosynthesis such as PsbS and LHCSR (Alboresi et al., 2010) were also similarly accumulated in the mutants as in WT plants (Figure S4).

**Table 1.** Pigment composition and PSII quantum efficiency of *P. patens* WT and *ndufb10* and *ndufa5* mutant lines. Chlorophyll (Chl) a/b ratio, Chl / carotenoids (car) ratio, total Chl content and PSII quantum yield (estimated by Fv/Fm) were evaluated in plants cultivated for 10 days in a minimal medium at 50 μmol photons m^-2^ s^-1^. For each measurement average ± SD (n ≥ 5) is reported.

| Genotype | Chl a/Chl b | Chl/Car | Chl content<br>($\mu\text{g mg}^{-1}\text{ DW}$) | Fv/Fm |
| --- | --- | --- | --- | --- |
| WT | $2.62 \pm 0.14$ | $3.73 \pm 0.37$ | $18.5 \pm 2.7$ | $0.79 \pm 0.01$ |
| *ndufb10* | $2.59 \pm 0.20$ | $3.65 \pm 0.40$ | $18.9 \pm 3.0$ | $0.78 \pm 0.02$ |
| *ndufa5* | $2.64 \pm 0.21$ | $3.60 \pm 0.42$ | $18.7 \pm 2.7$ | $0.79 \pm 0.01$ |

Photosynthetic functionality was analysed in more detail by fluorescence and NIR spectroscopy analyses. PSII maximum quantum yield (F_v_/F_m_) in dark-adapted plants was not altered by the mutations (Table 1). However, when plants were exposed to sub-saturating light conditions (330 μmol photons m^-2^ s^-1^) both *ndufa5* and *ndufb10* showed a higher yield for both PSI (Figure 5A) and PSII (Figure 5B), indicative of a higher electron transport rate. At the level of PSI, the higher yield was attributable to a lower donor side limitation (YND) while acceptor side limitation (YNA) was neglectable in WT as in mutant plants (Figure 5C-D). The lower PSI donor side limitation is consistent with a higher rates of electron transport to PSI and this is consistent with a less reduced quinone A (Q_A_) in the mutants than in the WT, as estimated by 1-q_L_ (Figure 5E). After actinic light was switched off, the estimated maximal quantum yield of PSI recovered to the same level in WT and mutants, whereas the recovery of PSII maximal yield and oxidation of Q_A_ was faster in the mutants. NPQ in *ndufa5* and *ndufb10* was activated as in WT plants but after a few minutes of illumination, it showed a partial relaxation in mutants (figure 5F).

**Figure 5.**
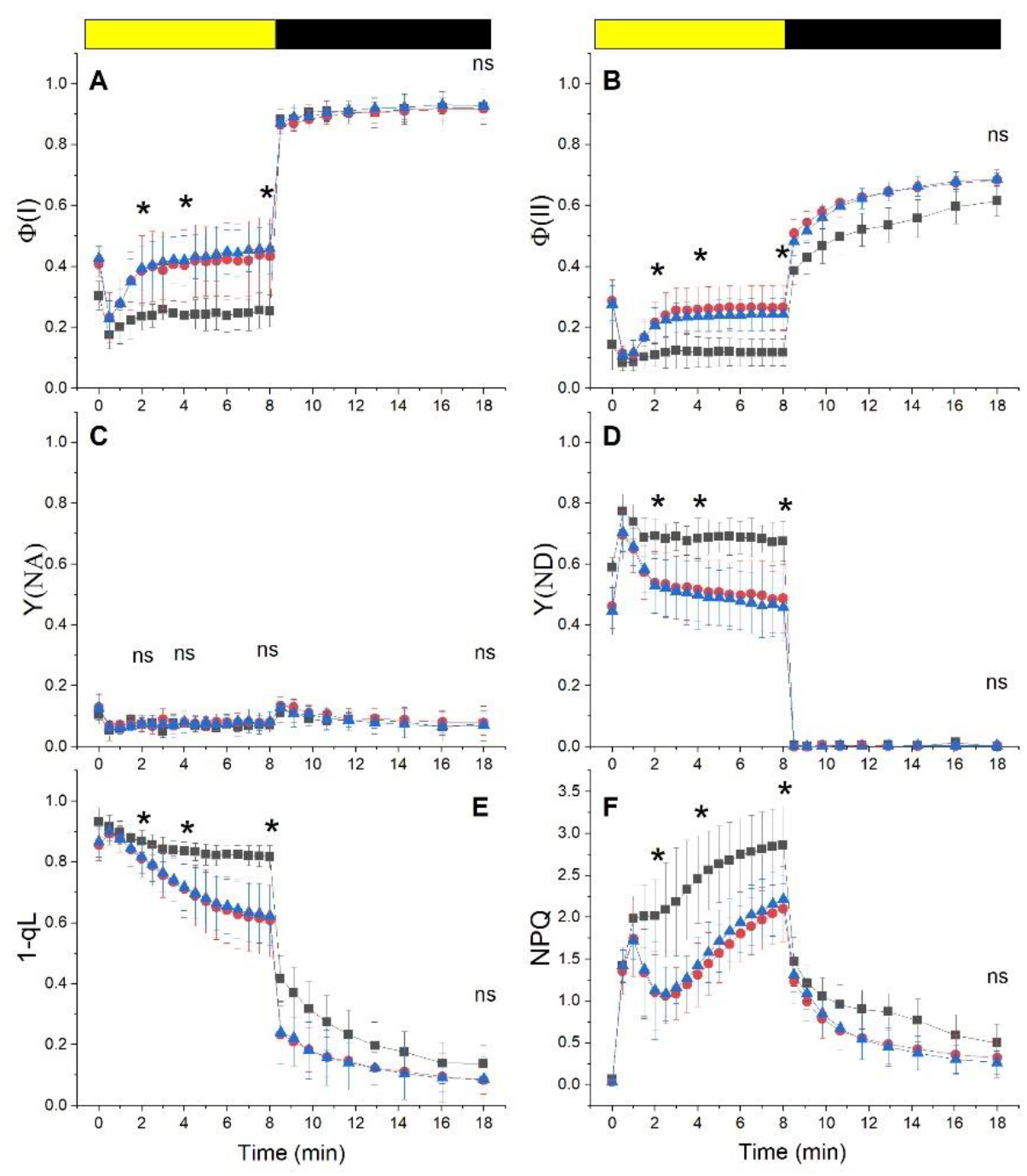
Alterations in Photosystem functionality in CI mutants. The yield of PSI (Φ(I), A), PSII (Φ(II), B). PSI acceptor side limitation (Y(NA), C), PSI donor side limitation (Y(ND), D). PQ redox state (1-q_L_; E) and non-photochemical quenching (NPQ; F) were measured with Dual PAM 100 in plants exposed to 330 μmol photons m^-2^ s^-1^ of actinic light intensity. Yellow / black bar indicates when actinic light was on / off respectively. WT, *ndufb10* and *ndufa5* KO are shown respectively with black squares, red circles and blue triangles. Data are shown as average ± SD (n > 4). Asterisks indicate statistically significant differences from WT plants (one-way ANOVA, n > 5, p < 0.01) after 2, 4 and 8 minutes of illumination and one after 10 minutes in the dark; ns indicates when eventual differences are not statically significant.

The same analyses showed similar results upon exposure to dim light (50 μmol photons m^-2^ s^-1^), corresponding to the illumination in the growth chamber (Figure S5). If plants were exposed to 2000 μmol photons m^-2^ s^-1^, a light intensity completely saturating photosynthetic capacity of the plants, WT and mutants showed instead no significant differences (Figure S6).

The impact of CI mutations on photosynthetic electron transport was further investigated by measuring electrochromic shift (ECS) caused by the generation of transmembrane potential at the level of the thylakoid membranes (Witt, 1979; Bailleul et al., 2010). The total ECS signal (ECS_t_), a proxy of the total proton motive force (pmf) generated, was found to be similar in WT and *ndufa5* and *ndufb10* mutants over a range of actinic light intensities of different duration (Figure 6A).

**Figure 6.**
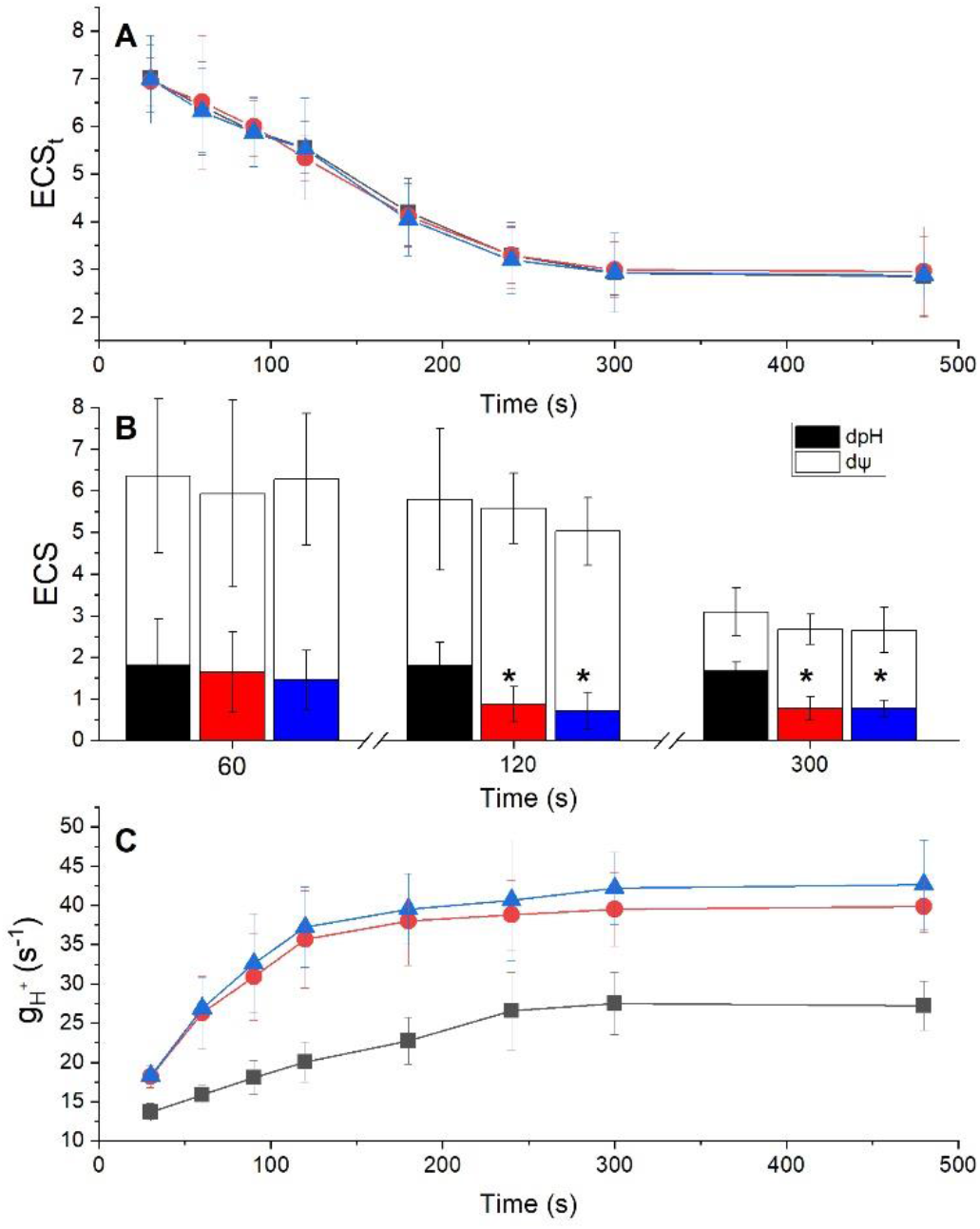
Impact of CI inactivation on photosynthetic electron transport and proton motive force. A) Total electrochromic signal (ECS_t_) generated by illumination with sub-saturating light (300 μmol photons m^-2^ s^-1^) at different intensity. ECS_t_ was quantified as (ECS_520_ - ECS_546_) / PSI Charge Separation and indicative of the proton motive force (pmf) generated. WT, *ndufb10* and *ndufa5* KO are shown respectively in black, red and blue. Average ± SD (n ≥ 5) is reported. B) quantification of total pmf as well as its partition in different components. ΔpH is shown in black, red and blue for WT, *ndufb10* and *ndufa5* KO respectively. The electric component (ΔΨ) in shown in white. Averages ± SD (n ≥ 5) are reported and asterisks indicate statistically significant differences (one-way ANOVA, n > 5, p < 0.01). D) g_H_^+^ is the proton conductivity of protons across the thylakoid membrane and it reflects the activity of ATPase. This was measured after exposing plants to light with different duration., WT, *ndufb10* and *ndufa5* KO are shown respectively as black squares, red circles and blue triangles.

ECS relaxation kinetics measured after the light was switched off after 300 seconds of illumination, however, indicated an altered partition between the electric and pH component of the transmembrane potential (Figure 6B) (Cruz et al., 2001). It is interesting to point out that this difference emerged as a consequence of the illumination since it became detectable only if plants were exposed to light for more than 120 seconds (Figure 6B and Figure S7). This suggests that the alterations in pmf partitioning emerged as a result of modifications in the photosynthetic activity (Avenson et al., 2005; Wang and Shikanai, 2019). This difference between genotypes was again light intensity-dependent and using a stronger, fully saturating, illumination (1000 μmol photons m^-2^ s^-1^) mutants were indistinguishable from WT (Figure S8).

To assess the impact of proton translocation, conductivity (g_H_^+^) from ATPase complex was estimated from the decay of ECS signal after the light was switched off. In both *ndufa5* and *ndufb10* the ECS decay kinetics were faster than WT, indicating that proton conductivity due to ATPase was higher (Figure S9). Monitoring of *g*_H_+ using actinic illumination of different duration (between 30 and 480 seconds) showed that *g*_H_+ in WT plants increased with the duration of the illumination reaching a steady state after approx. five minutes, consistently with a progressive activation of ATPase (Figure 6C), following a modulation of chloroplast ATPase activity by the sensing of stromal metabolic status during steady-state photosynthesis (Takizawa et al., 2008; Kohzuma et al., 2013). Interestingly, in *ndufa5* and *ndufb10* mutants the *g*_H_+ was higher but also more rapidly activated and it reached the maximal activity after about 120 seconds of illumination instead of the 300 seconds required by WT plants (Figure 6C).

Lumenal pH is a major signal for regulation of photosynthesis and its decrease under strong illumination is known to activate protective mechanisms such as heat dissipation of excess energy (non-photochemical quenching, NPQ) and the xanthophyll cycle (Li et al., 2009). Low lumenal pH is also known to inhibit cytochrome *b_6_f* activity to avoid over-reduction of PSI, a mechanism known as photosynthetic control (Nishio and Whitmarsh, 1993). When photosynthetic control is active, the electron transport rate from plastoquinol (PQH_2_) is slower, to reduce possibilities of PSI over-reduction. The impact of CI mutations on Cyt *b_6_f* activity was assessed by monitoring Cytochrome *f* (Cyt *f*) oxidation state from an absorption signal at 554 nm (Figure 7A, see methods for details). The oxidation state in illuminated samples can be estimated by comparing the signal with plants treated with PSII and Cyt *b_6_f* inhibitors DCMU and DBIMB where electron transport from PQ is blocked and thus Cyt f is fully oxidized. In *ndufb10* and *ndufa5* illuminated with sub-saturating light (300 μmol photons m^-2^ s^-1^) Cyt f is less oxidised than in WT plants (Figure 7B). When the light was switched off Cyt f reduction in mutants was also faster than in WT (Figure 7C), thus suggesting that in these plants electron transport from PQH_2_ is faster (Stiehl and Witt, 1969).

**Figure 7.**
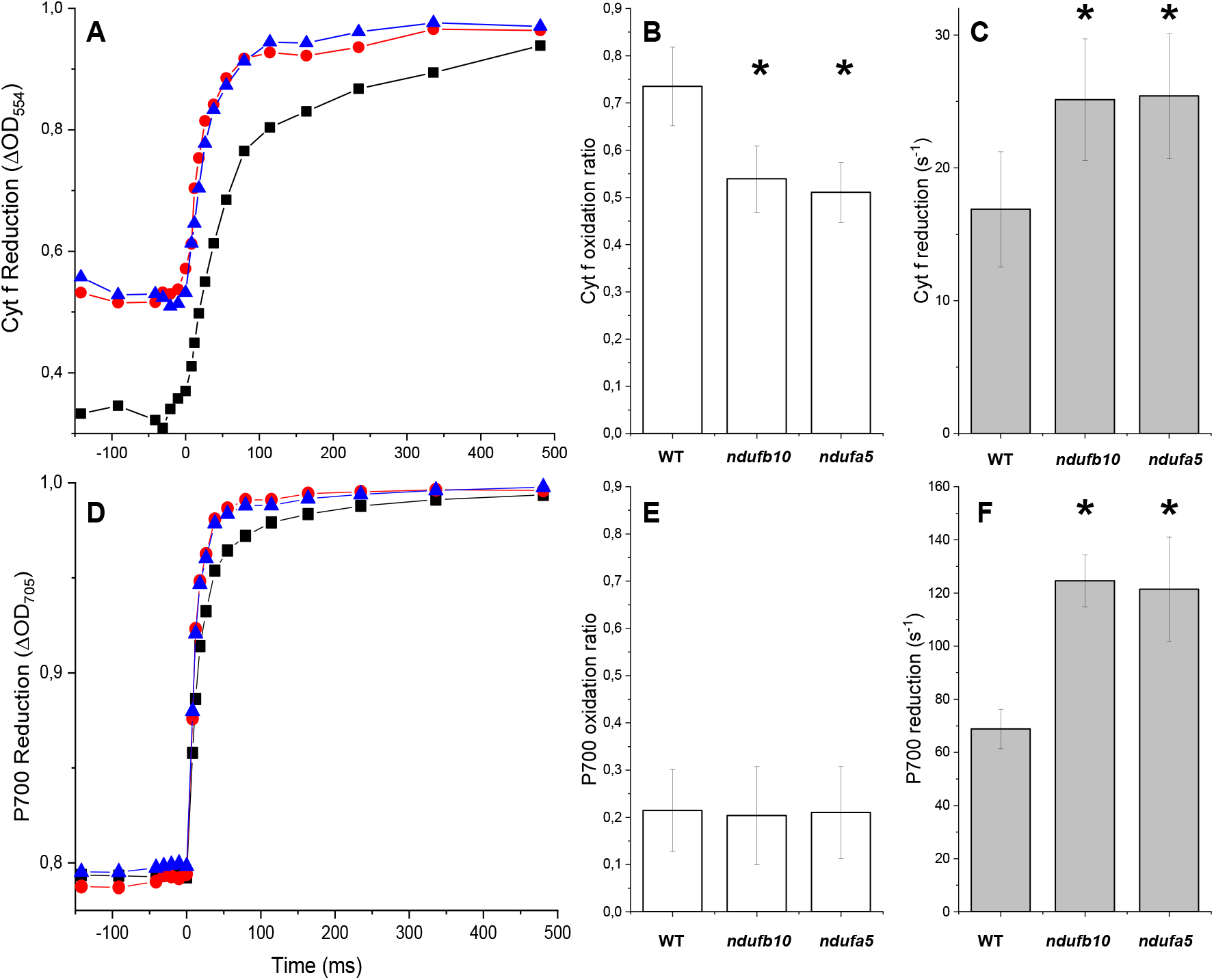
Cytochrome f and PSI oxidation. A) Cytochrome f oxidation monitored from ΔOD_554_ in dark-adapted plants subjected to constant illumination (300 μmol photons m^-2^ s^-1^) for 480 s before the light was switched off. B) Oxidation state is expressed as the ratio between Cyt f signals at the end of illumination and the maximal oxidation levels obtained by addition of DCMU/DBMIB to obtain complete oxidation. Reported values are average ± SD (n ≥ 4). C) Cyt f reduction rate in the different genotypes after 480 s actinic light exposure, quantified from half-time reduction. WT, *ndufb10* and *ndufa5* KO are shown respectively as black squares, red circles and blue triangles. Curves shown are the average of 4 independent measurements. For A and C asterisks indicate statistically significant differences (one-way ANOVA, n > 5, p < 0.01). D) PSI oxidation monitored from ΔOD_705_ in dark-adapted plants exposed to constant illumination (300 μmol photons m^-2^ s^-1^) for 480 s. B) P700 oxidized fraction is expressed as the ratio between P700^+^ signals at the end of illumination and the maximal oxidation levels obtained by addition of DCMU and DBMIB to block the PSI re-reduction. Reported values are average ± SD (n ≥ 4). F) P700^+^ reduction rate estimated from half-time reduction after 480 s actinic light exposure in the different genotypes. WT, *ndufb10* and *ndufa5* KO are shown respectively as black squares, red circles and blue triangles. For F asterisks indicate statistically significant differences (one-way ANOVA, n > 5, p < 0.01).

In the same measuring conditions, the impact on PSI was monitored from P700 oxidation state using a similar spectrophotometric approach, monitoring a differential absorption signal at 705 nm (ΔOD_705_, Figure 7D). CI mutants showed similar P700 oxidation level compared to WT plants at steady-state illumination (Figure 7E). However, when the light was switched off the rate of P700 reduction was faster in the mutants, suggesting that electrons flux toward P700 was more rapid (Figure 7F) consistent with a higher PQH_2_ oxidation rate.

## Discussion

### Inactivation of Complex I by targeting two different subunits yielded the same effects

In this work, independent KO lines targeting two subunits of CI were generated and verified to have the insertion of the resistance cassette in the expected genome position and to lose the mRNA expression (Figure 1 and Figure S1). *ndufa5* KO show complete depletion of CI activity in BN-PAGE (Figure 3B) and it did not accumulate NAD9 (Figure 3A), a core subunit of the Q module that is directly involved in the transfer of electrons to the ubiquinone which is required for the complex functionality (Massoz et al., 2013). This is consistent with previous results showing that NDUFA5 interacts with NAD7 and NAD9 to form an 80 KDa sub-complex that successively is integrated into the CI (Ligas et al., 2019). The second targeted subunit, NDUFB10, is instead integrated later during CI assembly and, in its absence, there is the formation of an incomplete CI of 800 kDa including both the N and Q modules but missing the proton pumping module and thus the biological activity (Barbieri et al., 2011). Despite this partial retention of some subunits (Figure 3), both mutants have inactive CI as shown by the insensitivity of oxygen consumption to the specific inhibitor rotenone (Figure 4B). In both lines, oxygen consumption in the dark is enhanced compared to WT plants. This can be explained by the re-organization of respiratory apparatus with increased CII and CIII content and a strong over-accumulation of AOX detectable both at the protein and activity level (Figure 3–4). Such reorganization of respiratory complexes allows compensating for the absence of CI activity, that in WT accounts for approx. 40% of O_2_ consumption rate in the dark (Figure 4B), reaching O_2_ evolution activities even higher than WT (Figure 4A). Such an enhanced oxygen consumption activity, however, is not expected to translate in a full recovery of the capacity of ATP biosynthesis since CII and alternative pathways like AOX do not contribute to the generation of transmembrane potential as CI (Vanlerberghe et al., 2020).

Alterations in respiratory complexes were also observed in CI mutants from *Arabidopsis* (Fromm et al., 2016b) and *Nicotiana* (Vidal et al., 2007) that also showed increased AOX accumulation and higher oxygen consumption in the dark. In *Arabidopsis*, it was shown that increased oxygen consumption in the dark is a distinctive feature of complete CI KOs, while this is not observed in CI partial mutants or KD lines (Kühn et al., 2015). Based on this observation, the increase in O_2_ consumption observed in *P. patens ndufa5* and *ndufb10* KO is a further confirmation that both lines have fully inactivated CI.

It is remarkable to observe that *ndufa5* and *ndufb10* KO show the same phenotypic differences with respect to WT in all analyses performed (Figure 2–8), even if the targeted genes were different. This similarity strongly suggests that plants responses observed are attributable to the lack of complex I activity, while the impact of specific subunits is minor, at least on the phenotypes analysed.

### Impact of respiration on photosynthesis adapted during evolution

All *P. patens* cells in various developmental stages contain chloroplasts and are photosynthetically active even if at different levels (Sakakibara et al., 2003). Here plants were analysed 10 days after inoculum on a mineral media containing no reduced carbon and were mainly composed by chloronema, a developmental stage containing phototrophic cells particularly rich in chloroplasts (Furt et al., 2012). The combined choice of the model organism and developmental stage, thus, should allow assessing the impact of respiration on photosynthesis without the influence of heterotrophic tissues like roots that are expected to be largely impacted by the lack of respiration. Despite the lack of heterotrophic cells, *P. patens* plants depleted in Complex I present a strong phenotype with a drastically impaired growth (Figure 2), demonstrating that respiration plays a major role in photosynthetically active cells in plants.

Plants exposed to 24 hours continuous light did not show any growth recovery suggesting that the impact of respiration is also not limited to sustaining metabolism during the night (Figure 2 and Figure S2). Even more strikingly, while WT plants exposed to continuous illumination show faster growth (Figure S2), this is not the case for CI mutants that are not able to exploit the extra light energy available. This show that inactivation of respiration directly impacts photosynthetic metabolism in *P. patens* plants and their ability to use light energy for growth.

The phenotypes of CI mutants in *P. patens* and *Arabidopsis* shows many similarities, such as the strong impact on growth and increased oxygen consumption activity, as the result of a re-organization of respiratory apparatus and an increased AOX content which ultimately lead to an increase of the electron flow through the alternative pathway with the purpose to sustain cellular ATP demand (Figure 3–4) (Kühn et al., 2015; Fromm et al., 2016a). This suggests that those phenotypes observed in *Arabidopsis* (Kühn et al., 2015) or tobacco (Vidal et al., 2007) are also attributable to effects on photosynthetic active cells, beyond the role in supporting metabolism during the night or in heterotrophic cells like roots.

CI mutants in the green alga *Chlamydomonas reinhardtii*, instead, have a growth rate close to WT in photoautotrophy and only show a decrease in duplication rate in heterotrophic conditions (Salinas et al., 2014; Larosa et al., 2018). Complex I mutants in *Chlamydomonas* also do not experience a reorganization of respiratory apparatus and showed a reduced oxygen consumption activity (Barbieri et al., 2011; Massoz et al., 2013; Larosa et al., 2018). Available information thus suggests that in plants respiration has a larger influence in photosynthetic metabolism than in *Chlamydomonas reinhardtii*.

While it is not possible to drive general conclusions on a highly diverse group such as eukaryotic algae-based only on information from a few species, these examples suggest the interaction between photosynthesis and respiration changed during evolution. Plants rely on photo-autotrophy and normally cannot complete their developmental cycle in the dark even if reduced carbon is supplied (Neff et al., 2000). The fact that obligate photo-autotrophic organisms rely more on respiration seems counterintuitive, but it can be explained considering that in these organisms mitochondrial respiration activity gradually adapted to work together with photosynthesis, optimizing the metabolite fluxes between organelles (Shameer et al., 2019). Respiration thus could gradually assume a major role in essential processes for photosynthesis such as redox balance and photorespiration, becoming itself essential for cells.

On the other hand, *Chlamydomonas* can growth in both autotrophic and heterotrophic conditions depending on the presence of carbon sources (Harris, 2001), an adaptation to a growing environment where light is not always present and cells must rely on other sources of energy to support their metabolism (Yang et al., 2015). *Chlamydomonas* cells can be exposed even to anaerobic conditions when it can activate fermentative metabolism and hydrogen production (Ghirardi et al., 2007; Grossman et al., 2011; Grechanik et al., 2020). In those anaerobic conditions, oxygen is absent and thus respiration is fully inactivated but still cells can perform photosynthesis (Godaux et al., 2015) that must thus be independent of respiratory activity. In organisms with such large metabolic flexibility and particularly if they are exposed to anaerobic conditions, respiration while important cannot assume an essential role in the photosynthesis, explaining why they are less impacted by a inactivation.

### Complex I Deficiency alters chloroplast ATP synthase activity and photosynthetic control of electron transport

Plants depleted in CI did not show any major alteration in the composition of the major photosynthetic pigments (Table 1) and complexes, PSI, PSII, Cyt *b6f* with only a slight increase in Chloroplast ATPase (Figure S4). Despite these similarities when plants are exposed to sub-saturating illumination (300 μmol photons m^-2^ s^-1^ and below) show several differences in photosynthetic functionality. Both PSI and PSII show a higher yield in mutants, suggesting a lower saturation level in illuminated plants and thus higher transport rate. CI mutants also show increased protein conductivity across the thylakoids membrane that, combined with an equivalent proton motive force, is again consistent with a higher electron transport rate.

The higher *g*_H_+ observed can be attributed to a higher ATPase activity. Chloroplast ATPase is in fact slightly over-accumulated in the mutants but it is also activated faster during the dark to light transitions in the mutants (Figure 6C and Figure S9) than in WT. ATPase activity is also inhibited by the depletion of inorganic phosphate (Pi) or ADP pool through metabolic feedback (Takizawa et al., 2008) and the higher activity in CI mutants could also be the result of a less effective metabolic control. The higher activity of chloroplast ATPase would thus be a response to satisfy an increased demand for ATP, compensating for a reduced contribution from mitochondrial respiration.

The higher ATPase activity causes a faster translocation of protons in the stroma and an increase in the lumenal pH (Figure 6B), in turn, affecting photosynthesis regulatory mechanisms modulated by lumen acidification. The WT and mutant lines accumulated similar levels of NPQ activators PsbS or LHCSR (Figure S4) and, consistently, the maximal NPQ capacity is equivalent in all genotypes (Figure S6F). However, mutants exposed to sub-saturating illumination show a relaxation of NPQ after a few minutes (Figure 5F) that can be explained by a reduction of ΔpH across the thylakoids membrane ((Joliot and Finazzi, 2010), Figure 6B-C).

An altered lumen acidification is also expected to affect photosynthetic control, the modulation of PQH_2_ oxidation at the Cyt *b6f*. Mutants exposed to sub saturating illumination indeed showed a faster rate of Cyt *f* reduction (Figure 8C), suggesting a higher electron transport activity and a lower photosynthetic control. This hypothesis is consistent with several other differences observed such as the higher efficiency in both photosystems (Figure 5A-B and Figure S5A-B), a less pronounced PSII acceptor limitation (1-q_L_) (Figure 5E) and PSI donor limitation (YND) (Figure 5D and Figure S5D) compared to WT. PSI also shows faster reduction kinetics (Figure 8F). All the differences observed between the WT and mutants, summarized in Figure S10, can be explained with a decreased photosynthetic control and thus a higher Cyt *b_6_f* activity, thus increasing the rate of transport of electrons from PQ to PSI as the result of higher ATPase activity and higher lumenal pH (Takizawa et al., 2007; Takizawa et al., 2008).

It is interesting to observe that when light is in strong excess (1000-2000 μmol m^-2^ s^-1^, Figure S6 and Figure S8), thus 25-50 times higher than the illumination used for plants growth, the mutants become indistinguishable from WT. This observation suggests that with a very strong illumination the eventual differences in ATPase conductivity are uninfluential, an expected behaviour if in these conditions the downstream metabolic reactions became strongly saturated. This further confirms the conclusion that photosynthetic apparatus is highly similar in WT and CI mutants and that the phenotypes observed in the latter are due to the depletion of respiration that affects the regulation of photosynthesis within the chloroplast. This could be explained by idea that ATP pools in chloroplast, mitochondria and cytosol are exchangeable, if not directly through transport of reduced molecules that are used for ATP biosynthesis (Shameer et al., 2019) and thus if ATP biosynthesis is reduced in the mitochondria because of CI inactivation it causes indirect effects on photosynthetic activity.

## Acknowledgments

We thank Dr. Etienne Meyer (Institute of Plant Physiology; the Martin-Luther-University, Halle-Wittemberg) for providing the NAD9 antibody and Prof. Hans-Peter Braun (Institute of Plant Genetics; Leibniz University, Hannover) for the SDH1-1, α-MPP, COX2 and ATPase β-subunit antibodies.

## Materials and Methods

### Plant material and growth. P. patens

*P. patens* (Gransden) Wild-Type (WT), *ndufb10* and *ndufa5* KO lines were cultivated in the protonemal phase by vegetative propagation on PPNH_4_ medium (Ashton et al., 1979) and grown under standard conditions: 24°C, long photoperiod (16:8 light: dark) with 50 μmol photons m^-2^ s^-1^. Physiological and biochemical characterizations were performed on 10 day-old tissue cultivated in PPNO_3_ medium (Ashton et al., 1979). The growth rate in all the media and light conditions was evaluated starting from protonema colonies of 2 mm in diameter followed for 21 days. Colony size was measured as in Storti et al., (2019). In brief, high-quality images (600 ppi) were acquired using a Konica Minolta Bizhub C280 scanner. Images were analysed with FIJI (https://fiji.sc/) using the ‘threshold colour’ plugin to subtract the plate background. Colony size was calculated from the integrated density (area x mean density). This strategy was chosen to take into account the development from 2D (chloronema and caulonema) to 3D structures (gametophore and rhizoids) of moss colonies which are lost considering only the area.

### Moss transformation and mutant selection

The *ndufb10* and *ndufa5* KO constructs were employed to mutate the *Ndufb10* gene and *Ndufa5* gene respectively (Figure S1). The transformation was performed through protoplast DNA uptake as described in Alboresi et al. (2010). 6-day old protonema cells growth in PpNH_4_ medium were treated with fungal driselase (Sigma-Aldrich) to break the cell wall. Resulting protoplasts were filtered with 100 μm micro cloth. Protoplasts were washed and resuspended in PEG-4000-containing solution, mixed with digested linear DNA from KO constructs (20 μg) and exposed to heat shock (5 min at 45°C) to open cell membrane. After 1 day of recovery, protoplasts were first immersed in a top layer solution and plated on agarified medium added with mannitol to prevent their lysis. After 7 days, recovered cells were moved to a new plate with antibiotics for transformants selection. The selection was repeated twice. Resulting transformants were homogenized using 3 mm zirconium glass beads (Sigma-Aldrich), and genomic DNA (gDNA) was isolated according to a rapid extraction protocol (Edwards et al., 1991) with minor modification. PCR amplifications of recombination cassette were performed on extracted gDNA (Table S1; Fig. 1, Fig. S1). To confirm that *ndufb10* and *ndufa5* KO lines were lacking target gene expression Reverse-Transcriptase PCR (RT-PCR) was performed on cDNA (RevertAid Reverse Transcriptase; Thermo Scientific) synthesized after RNA extraction.

### Western blot analysis

Total protein extracts were obtained by pestled protonema tissue in sample buffer (50 mM TRIS pH 6.8, 100 mM DTT, 2% SDS and 10% glycerol). Samples total chlorophyll was quantified, and every well was loaded accordingly to the quantification. After SDS-PAGE, proteins were transferred to a nitrocellulose membrane (Pall Corporation). Membranes were hybridized with specific primary antibodies: anti-PsaA, Agrisera, catalogue number AS06172; anti-Cyt *f*, Agrisera, catalogue number AS06119; anti-γ ATPase, Agrisera, catalogue number AS08312, anti-AOX1/2, Agrisera, catalogue number AS04054; custom made anti-NAD9 (Schimmeyer et al., 2016); custom made anti-SDH1-1 (Complex II),anti-α-MPP (Complex III), anti-COX2 (Complex IV), anti-β subunit (Complex V) (Peters et al., 2012) and custom made anti-D2, anti-PSBS and anti-LHCSR (Storti et al., 2019). After hybridization, signals were detected with alkaline phosphatase-conjugated antibody (Sigma Aldrich).

### Crude membrane extracts preparation

Crude membrane extracts were prepared as in Pineau et al., (2008) with minor modifications. Approximately 300 mg of fresh or frozen (−80 °C) protonema grown in PpNO3 for 10 days were homogenized in 2 ml of 75mMMOPS-KOH, pH 7.6, 0.6Msucrose, 4mM EDTA, 0.2% polyvinylpyrrolidone 40, 8 mM cysteine, 0.2% bovine serum albumin using a plant Potter glass tissue grinder pound at 0 °C. The homogenate was filtrated across one layer of micro cloth with 20 μm pores and centrifuged at 4°C at 1300 g for 4 min. The supernatant was collected and centrifuged again at 22,000 g for 20 min. The resultant pellet, which contained most of the thylakoid and mitochondria membranes, was resuspended in 200 μL of 10 mM MOPS-KOH, pH 7.2, 0.3 M sucrose. The protein concentration on crude membrane extracts was quantified using the BCA protein assay.

### Blue native protein electrophoresis (BN-PAGE)

Gels were cast in 8 × 10 cm plates using the buffer described by Kügler et al., (1997) with an acrylamide gradient of 4–12% in the running gel and 4% acrylamide in the stacking gel. A volume of crude membrane extracts corresponding to 100 mg of proteins wash washed with three volumes of H2O Milli-Q^®^ and centrifuged at 4 °C, 21,470 g for 20 min. The pellet was resuspended in 20 μL of ACA buffer 1x (50 mM Bis-Tris, pH 7.0; 750 mM aminocaproic acid; 1 mM EDTA) Järvi et al., (2011). For protein solubilization 20 μL of β-dodecyl maltoside (β-DM) 2% (w/v) prepared in ACA buffer were added to the tube, to reach a final volume of 40 μL, reaching β-DM 1% in ACA buffer. Each tube was vortexed for 30 seconds, kept on ice for 5 min and centrifuged at 4°C, 22,000 g for 8 min. The supernatant was supplemented with 4 μL of Coomassie Blue 5% solution (20 mM Bis-Tris, 500 mM aminocaproic acid, Coomassie Blue G-250 5 % (w/v)). Anode and cathode buffers were the same used by Järvi et al. (2011) for BN gel; the cathode buffer was supplemented with Coomassie Blue G-250 0.02 % (w/v). The gel was run at 75 V for 30 minutes. Then, the cathode buffer was replaced with fresh cathode buffer without Coomassie Blue and the gel was run at 100 V for 30 min, at 125 V for 30 min, at 150 V for 60 min, at 175 V for 30 min and at 200 V for 60 min. The total running time was about 4 hours.

### Determination of NAH dehydrogenase in-gel activity

After BN-PAGE, the NADH dehydrogenase activity of CI was revealed by incubation of the gel in the presence of 1 mM nitro blue tetrazolium (NBT) and 0.2 mM NADH in 50 mM potassium phosphate buffer (pH 7.0) (Barbieri et al., 2011).

### Oxygen consumption and oxygen evolution

Oxygen consumption and evolution were evaluated as in Storti et al. (2020) with a Clark-type O_2_ electrode (Hansatech, King’s Lynn, UK). In brief, two protonema disks of about 1 cm of diameter coming from 10 days old plates were introduced in the measurement chamber filled with 2 mL of a solution containing NaCO_3_ 0.1 mM maintained at 23°C. After ten minutes in the dark (respiratory rate), the light was turned on and the oxygen variation was recorded for other ten minutes (photosynthetic rate). O_2_ consumption and evolution rates were normalized to the total Chl content of each sample. Chl content was evaluated after extraction with 80% acetone (Porra et al., 1989). Inhibitors employed for respiratory analysis were 50 μM rotenone (Complex I), 1 mM KCN (Complex IV) and 2 mM SHAM (AOX). For inhibitor treatments, protonema tissue was incubated 30 minutes in the dark in medium supplemented with the inhibitor. Inhibitors were also added at the measuring chamber during oxygen consumption evaluation.

### Pigment analysis

10 days old protonemal cells were broken with a plastic grinder and pigments were extracted with 80% acetone. Whole-plant extracts in acetone were fitted with those of individual purified pigments to calculate Chl*a*/*b* and Chl/carotenoid (Chl/Car) ratios (Croce et al., 2002).

### Spectroscopic analyses

*In vivo* chlorophyll fluorescence and Near InfraRed (NIR) absorption analyses were performed at room temperature with a Dual-PAM 100 system (Walz) on protonema grown for 10 days in PpNO_3_ in WT and mutant lines. Before the analysis, the plants were adapted in the dark for 40 min, and the F_v_/F_m_ parameter was calculated as (F_m_-F_o_)/F_m_. Induction curves were obtained setting actinic red light at (approx.) 50, 330 or 2000 μmol photons m^-2^s^-1^, and photosynthetic parameters were recorded every 30 s. At each step, the photosynthetic parameters were calculated as follows: Φ(II) as (F_m_’-F_o_)/F_m_’, q_L_ as (F_m_’-F)/(F_m_’-F_o_’) x F_o_’/F and NPQ as (F_m_-F_m_’)/F_m_’, Φ(I) as 1-Y(ND)-Y(NA); Y(NA) as (P_m_-P_m_’)/P_m_; Y(ND) as (P - P_o_/P_m_) (Klughammer and Schreiber, 1994). Electrochromic shift (ECS) spectra were recorded with a JTS-10 system (Biologic) in plants that were dark acclimated for 40 min and imbibed with 20 mM HEPES, pH 7.5. and 10 mM KCl. For each measure, the background signal at 546 nm was subtracted from the 520 nm signal, in this way eliminating the contribution of scattering and cytochromes. Functional total Photosystems (PSs) quantification was performed by a single flash turnover using a xenon lamp. The light produced by xenon gas can induce PSI double charge separation, and thus PSI content could be overestimated by c. 40% but it does not affect the comparison of different samples (Gerotto et al., 2016). Moreover, samples were incubated with 20 μM 3-(3,4-dichlorophenyl)-1,1-dimethylurea (DCMU) and 4 mM HA (hydroxylamine) to calculate the contribution of PSI alone (Joliot et al., 2004). Total Electron Transport Rate (ETR) was measured with DIRK (dark-induced relaxation kinetic (Sacksteder and Kramer, 2000)) analysis as in Gerotto et al. (2016). Briefly, throughout 5 min of continuous illumination (at 350 or 940 μmol photons m^-2^ s^-1^, 630 nm LED) light was switched off at different times (0.24, 0.52, 0.79, 1.07, 1.52, 2.47, 3.93, 6.38, 8.83, 18.88, 48.9, 168.97, 288.99 s) for 20ms to follow fast relaxation of the ECS components (Slope in the Dark, SD). ETR was calculated as the ECS signal in light (SL) subtracted from the SD and normalized to the sample PSI content. At the end of the fifth min, the light was switched off to follow relaxation kinetics and evaluate the proton motive force (pmf) generated during light treatment as in Storti et al., (2020). The pmf was determined as the difference of the maximum signal at the light steady-state and minimum level of ECS in the dark and normalized to total charge separation PSI. The pmf portioning in ΔΨ and ΔpH was performed as in Baker et al., (2007). The ECS relaxation was followed for 30 s after light to dark transition and the ECS signal in this relaxed state was recorded. Difference between the relaxed ECS signal and the maximal signal at light steady-state is defined as ECS_ss_, which is proportional to ΔΨ. ΔpH was calculated from the difference between the relaxed ECS signal and a minimum level of ECS in the dark. Proton conductance (g_H_^+^) was estimated by fitting the first 300 ms of the ECS decay curve with a first-order exponential decay kinetic as the inverse of the decay time constant as described earlier (Avenson et al., 2005). g_H_^+^ was calculated after exposure to 30s, 60s, 90s, 120s, 180s, 240s, 300s and 480s of illumination (350 μmol photons m^-2^ s^-1^). P700^+^ absorption kinetics were calculated from the absorption at 705 nm (Δ_OD_705) and following the re-reduction after plants were illuminated for 5 min at 350 μmol photons m^-2^ s^-1^. Oxidized P700 was calculated by comparing the maximum signal from P700^+^ obtained before and after incubating plants with20 μM DCMU and 150 μM 2,5-Dibromo-6-isopropyl-3-methyl-1,4-benzoquinone (DBIMB). P700 kinetic rate was calculation by relaxation time t1/2 of P700^+^ reduction after light was switched off. Cyt *f* absorption was calculated as for P700^+^, except for the interference filter at 554 nm. In this case, 546 nm and 573 nm were used as background and removed to 554 nm signal (Finazzi et al., 1997). Oxidation status and Cyt *f* kinetic rate measurements were performed as for P700 measurements.

## Notes

### Competing Interest Statement

The authors have declared no competing interest.

