## Supplementary Material for "Inactivation of mitochondrial Complex I stimulates chloroplast ATPase in Physcomitrella (*Physcomitrium patens*)"

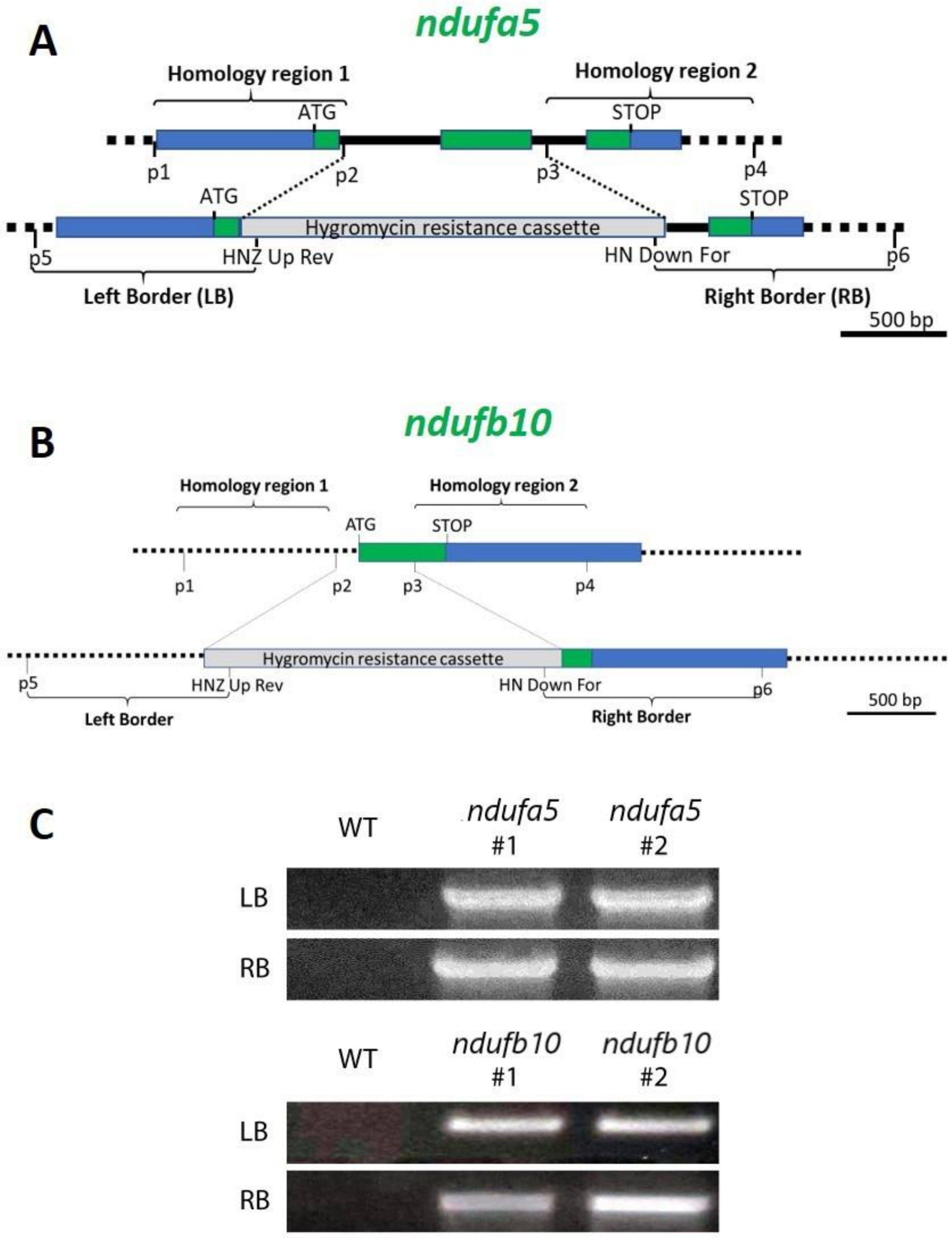


Figure S1. Generation of *P. patens* mutant lines lacking respiratory complex I A-B) KO scheme for *Ndufa5* and *Ndufb10* respectively, showing the CDS, the regions of homology chosen to drive homologous recombination and insertion of the resistance cassette. C) Example of PCR for verification of the homologous recombination event. PCR products (called Left and Right Border, LB and RB) are generated only if the resistance cassette is inserted in the expected genomic region.

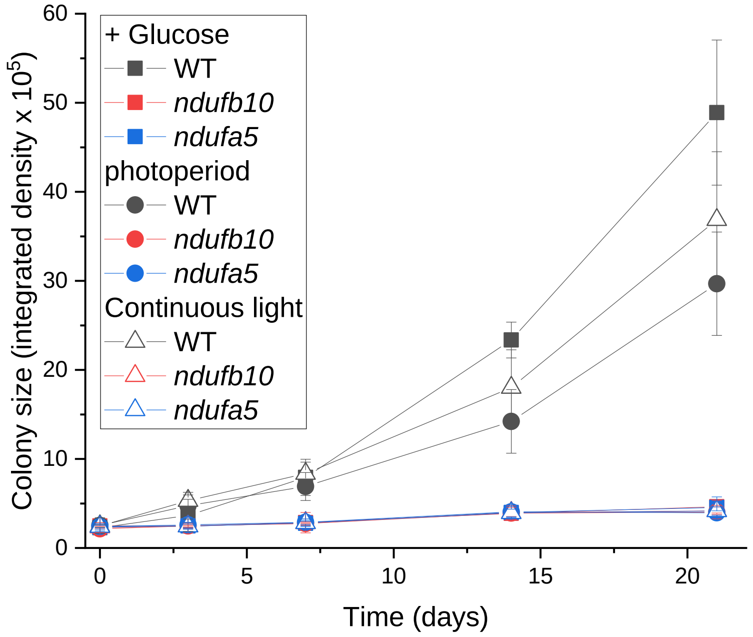


**Figure S2. Growth kinetics of *Physcomitrella patens* plants under different conditions.** Plants were grown under different light regimes and different medium composition as shown in Figure 2 and their size was evaluated after 0, 3, 7, 14 and 21 days of growth. All plants were exposed to 50 µmol photons $m^{-2}$ $s^{-1}$. Squares show growth in medium supplemented with 0.5% of glucose, circles growth with 16h light/8h dark photoperiod in minimal medium, triangles in continuous light regime in minimal medium. WT, *ndufb10* and *ndufa5* KO are shown respectively in black, red and blue. Average ± SD (n ≥ 5) is reported.

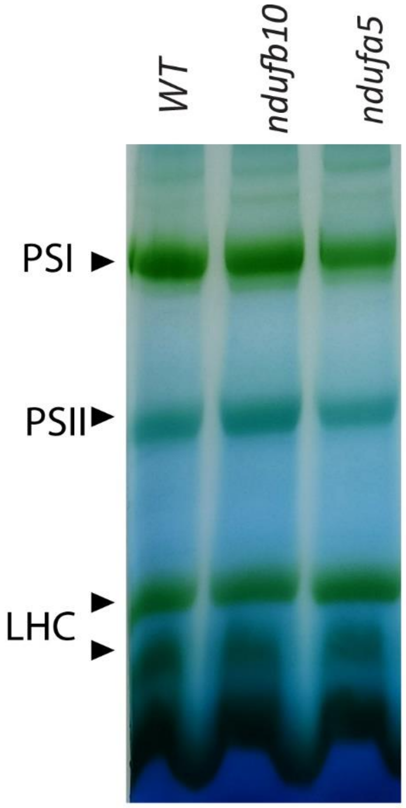


**Figure S3. BN-PAGE separation of crude organelles extracts.** Crude Membrane proteins extract from WT, *ndufb10* and *ndufa5* were solubilized with 1% of β-dodecyl maltoside (β-DM). Complexes of thylakoid membranes identifiable as green bands are indicated; PSI, Photosystem I; PSII, photosystem II; LHC, light-harvesting chlorophyll complexes.

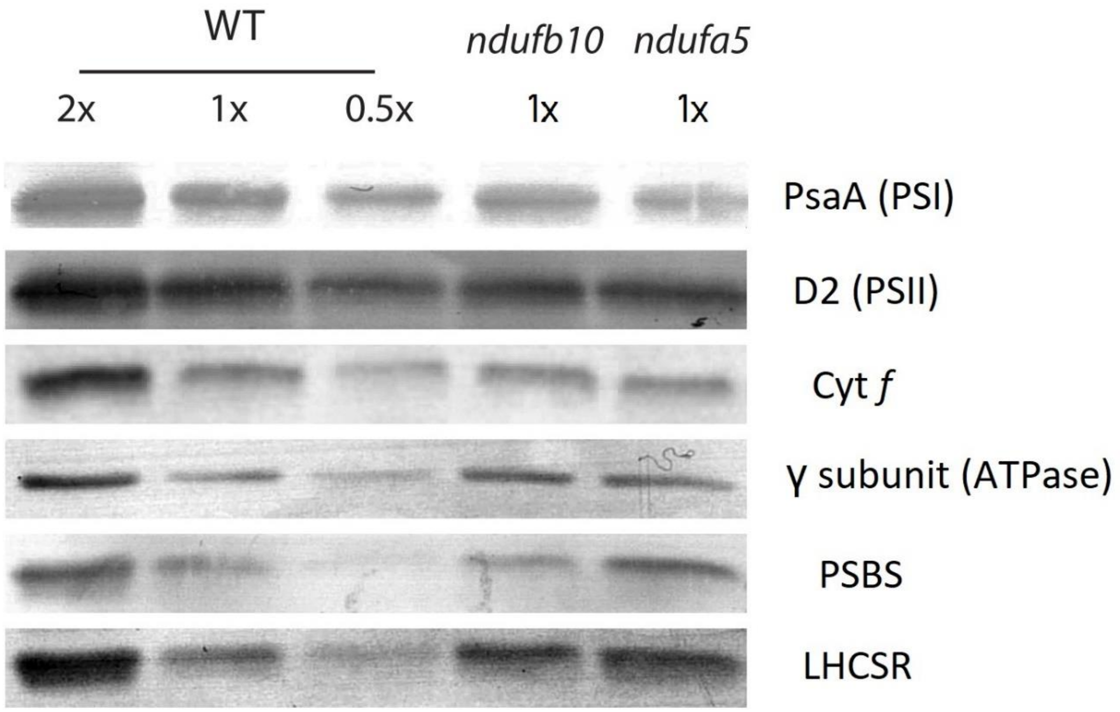


Figure S4. Impact of Complex I mutations on the photosynthetic apparatus composition. Immunoblot analysis of various proteins of the photosynthetic apparatus. For WT 1X, *ndufb10* and *ndufa5* a total protein extract amount equivalent to 2 µg of total chlorophyll was loaded.

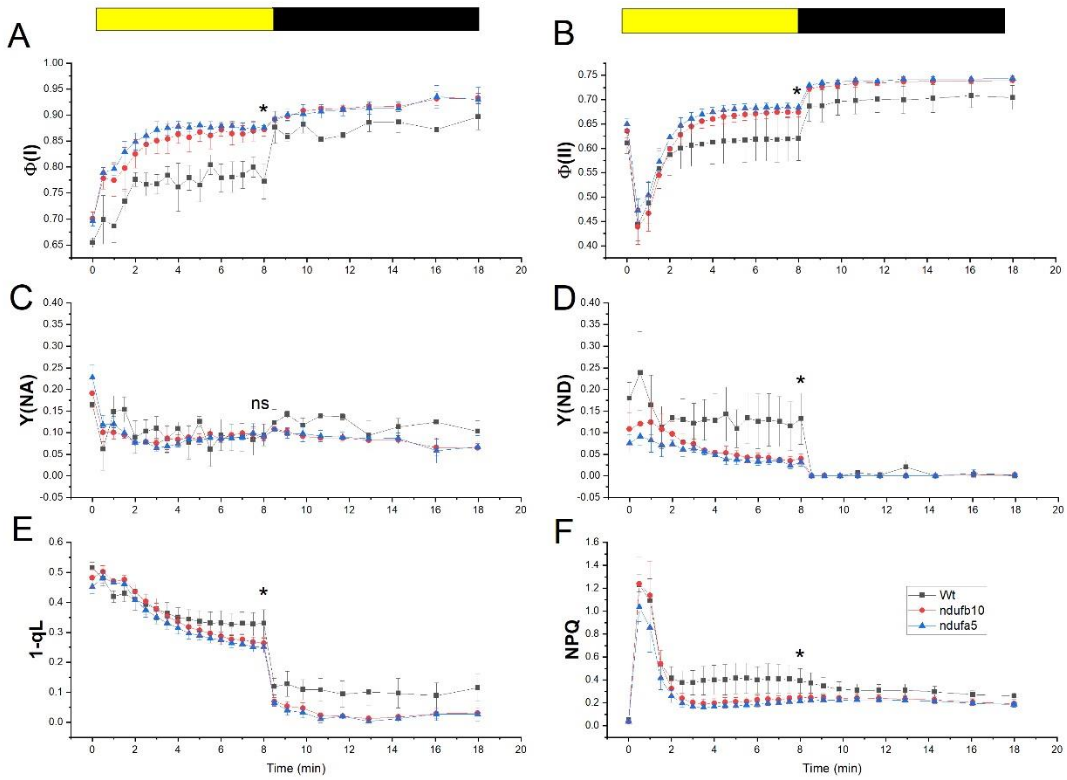


**Figure S5. Photosynthetic efficiency under dim illumination.** The yield of PSI (Φ(I), A), PSII (Φ(II), B). PSI acceptor side limitation (Y(NA), C), PSI donor side limitation (Y(ND), D). PQ redox state (1-qL; E) and non-photochemical quenching (NPQ; F) were monitored under illumination of 50 µmol photons $m^{-2}s^{-1,}$ corresponding to light intensity during growth. WT, *ndufb10* and *ndufa5* KO are shown respectively as black squares, red circles and blue triangles. Yellow/black bar indicates light on/off. Data are shown as average ± SD (n > 4). Asterisks indicate statistically significant differences (one-way ANOVA, n > 5, p < 0.01) between WT and both mutants while ns indicates when eventual differences are not statically significant. Statistical analyses were performed for the last point before light was switched off.

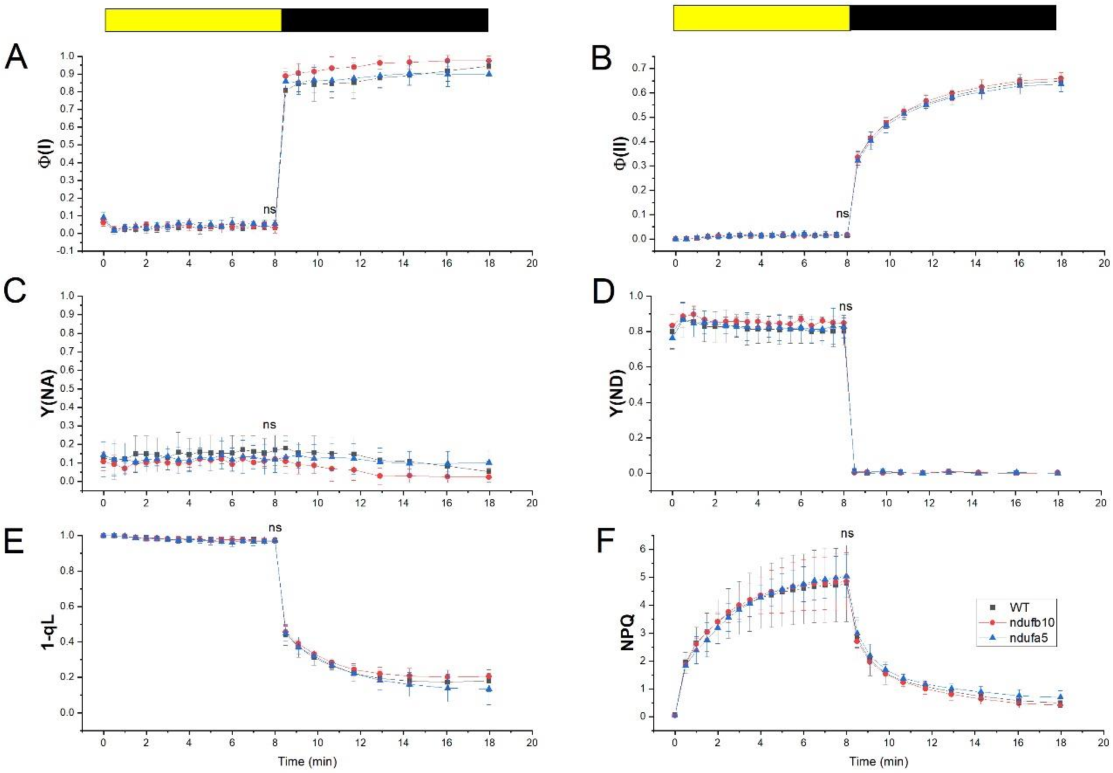


**Figure S6. Photosynthetic efficiency under saturating illumination.** The yield of PSI (Φ(I), A), PSII (Φ(II), B). PSI acceptor side limitation (Y(NA), C), PSI donor side limitation (Y(ND), D). PQ redox state (1-qL; E) and non-photochemical quenching (NPQ; F) under 2000 µmol photons $m^{-2}s^{-1}$ of light intensity. WT, *ndufb10* and *ndufa5* KO are shown respectively with black square, red circle and cyan triangle. Yellow boxes above the panels represent actinic light on, instead black boxes represent actinic light off. Data are shown as average ± SD (n > 4). Statistical analysis was performed for the last point of illumination before light was switched off. (ns = not significant).

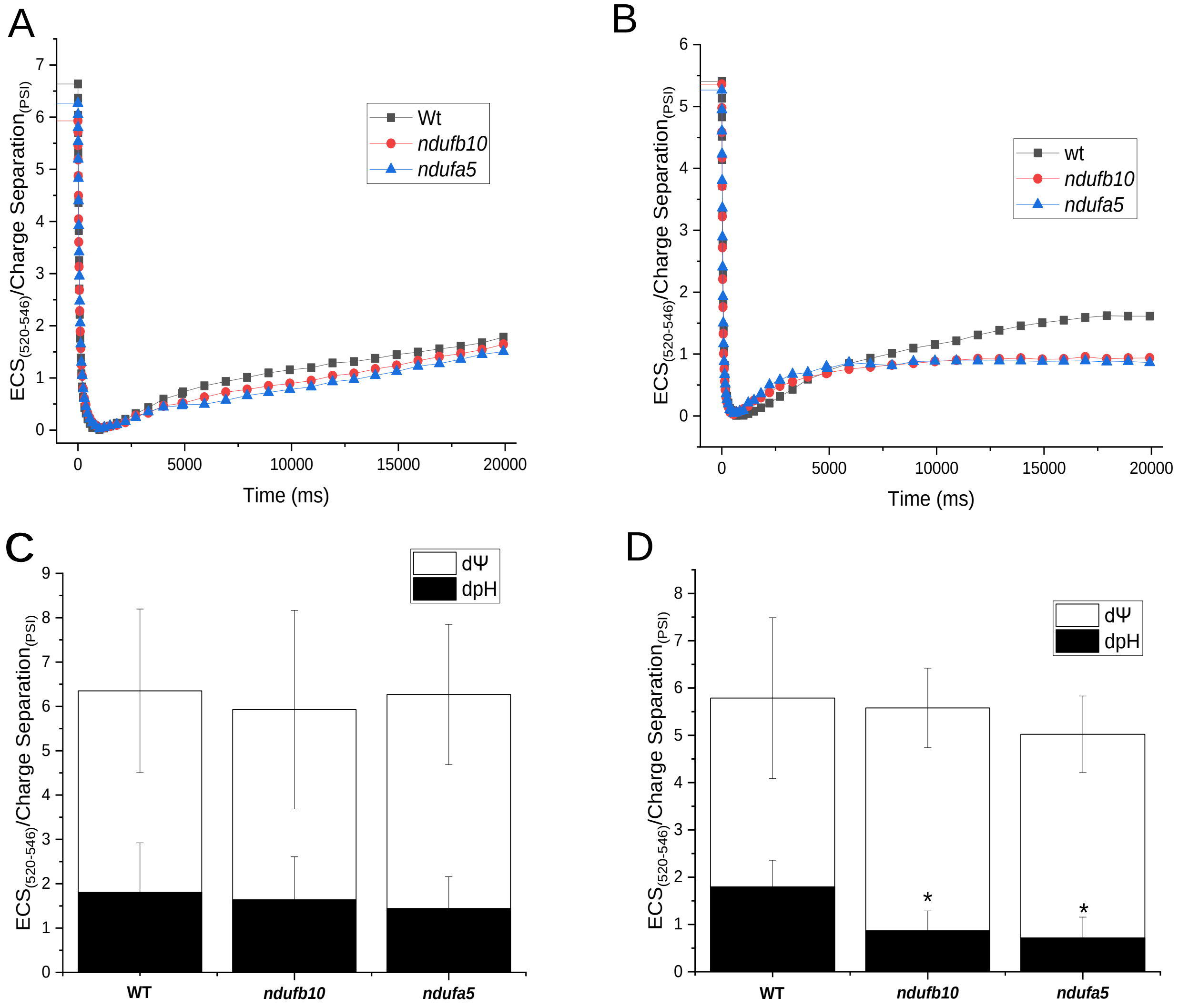


**Figure S7: Proton motive force composition before the steady state.** A-B) Representative traces of the ECS signal after the light is switch off after 60 s (A) and 120 s (B) of illumination (300 µmol photons $m^{-2}$ $s^{-1}$). WT, *ndufb10* and *ndufa5* KO are shown respectively as black squares, red circles and blue triangles. C-D) pmf partitioning after 60 s (C) and 120 s (D) is represented with orange columns for ΔpH component and with green columns for the electrical potential (ΔΨ). For D asterisks indicate statistically significant differences (one-way ANOVA, $n > 5$, $p < 0.01$).

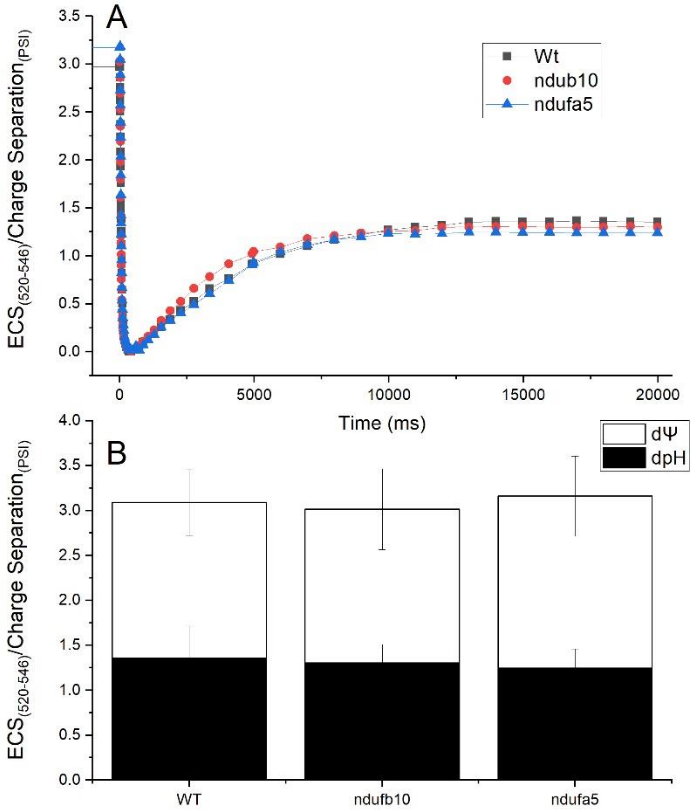


**Figure S8: Photosynthetic proton motive force at saturating illumination.** Proton motive force (pmf) generated in thylakoids membranes assessed by electrochromic signal in plants exposed for 300 seconds of illumination with saturating light (900 µmol $m^{-2}$ $s^{-1}$). A) Representative tracks of the ECS signal after light is switch off. WT, *ndufb10* and *ndufa5* KO are shown respectively with black square, red circle and cyan triangle. C) Black columns are representative for the osmotic components (ΔpH) of pmf while white columns show the electrical potential (ΔΨ).

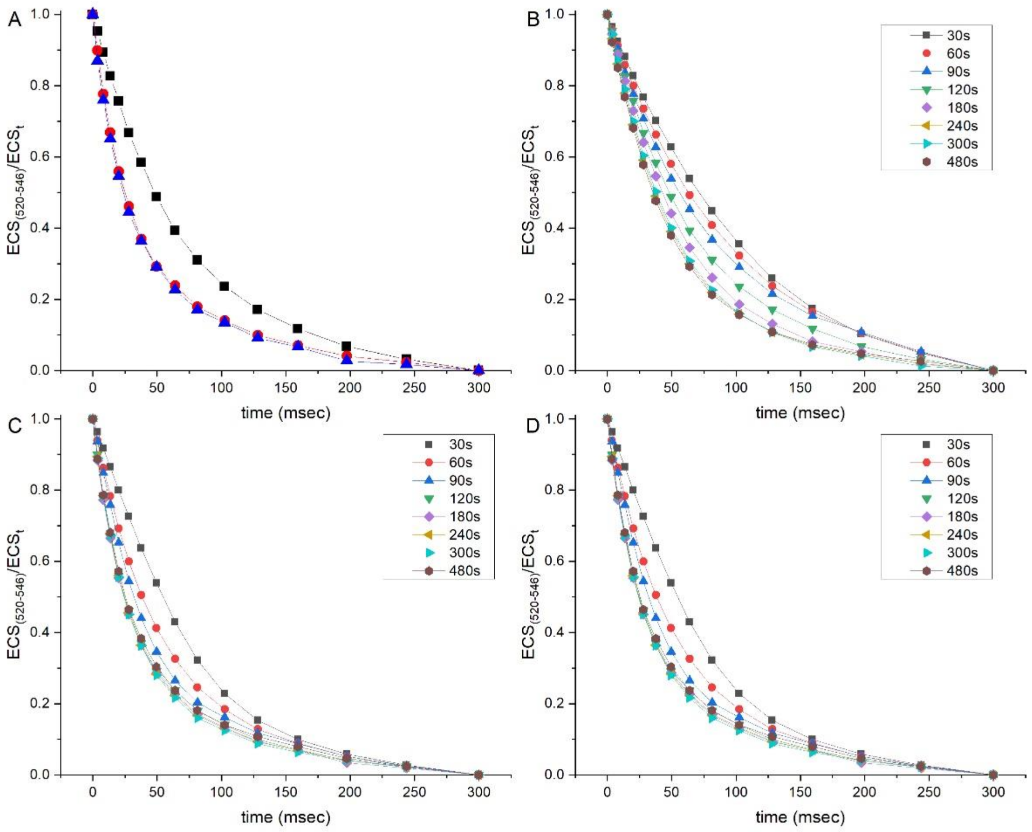


**Figure S9. ATPase activity assessed from ECS relaxation.** A) ECS relaxation kinetics after 120 seconds of illumination with sub-saturating light (300 µmol $m^{-2}$ $s^{-1}$). WT, *ndufb10* and *ndufa5* KO are shown respectively as black squares, red circles and blue triangles. B-D) ECS relaxation traces for WT (B), *ndufb10* (C) and *ndufa5* (D) measured after illumination of different length. In every panel traces after 30, 60, 90, 120, 180, 240, 300 and 480 seconds are shown respectively as black squares, red circles, blue triangles, green triangles, magenta squares, ochre triangles, cyan triangles and carmine hexagons.

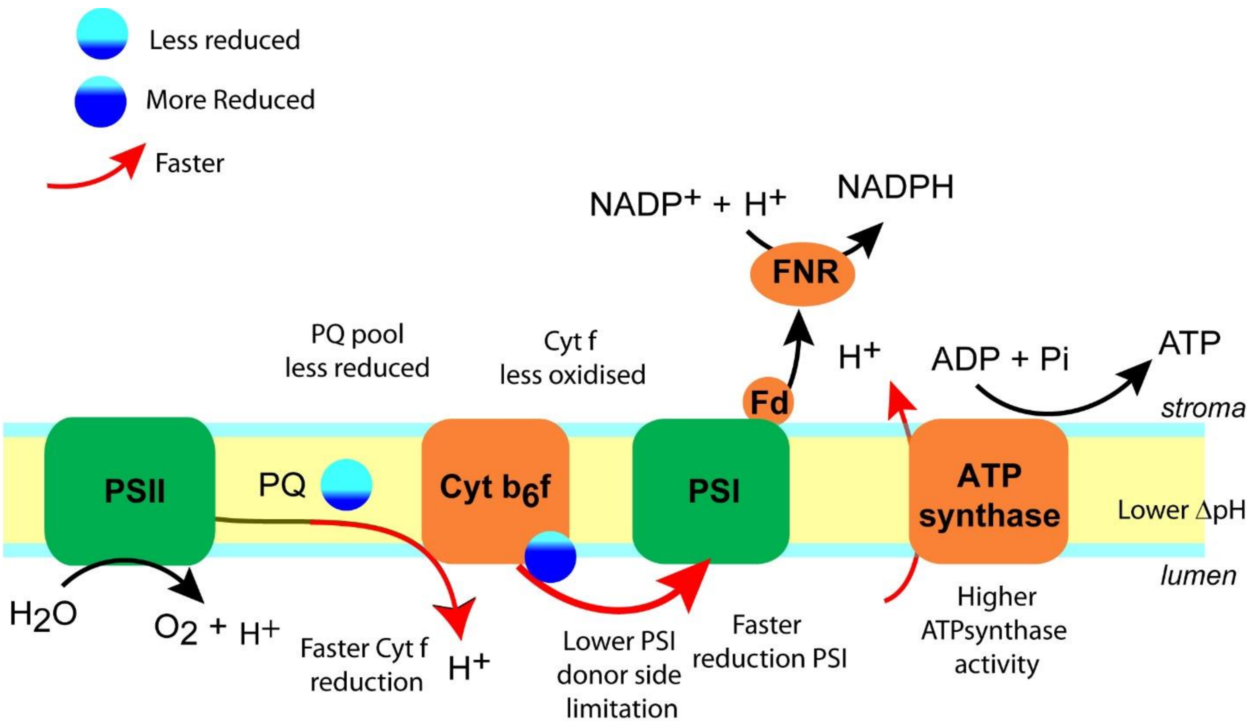


Figure S10. Summary of photosynthetic electron transport alterations in CI depleted plants.

**Table S1: Primers employed in this work**

| Gene | Primer Name | Sequence | Use |
|---|---|---|---|
| Ndufb10 | Ndufb10-p1 | GTTTAAACGGCCCTAATGACATAAGTCCC | KO isolation |
| | Ndufb10-p2 | CTCGAGGTGGATGAATGATGCTGGTG | KO isolation |
| | Ndufb10-p3 | ATCGATCAAGTCGAGAAGGCGAAGATC | KO isolation |
| | Ndufb10-p4 | TTAATTAAGATGGCGTCGAGTTCCTTCATC | KO isolation |
| | Ndufb10-p5 | GGATCCACAATCAGGAATTA | KO screening |
| | Ndufb10-p6 | ACCAAAGCAGTCCTGAAC | KO screening |
| | Ndufb10-RTf | GTTCAACAGCGACTACCCTA | RT-PCR |
| | Ndufb10-RTr | GAAACCCTTGTTCTGAAATG | RT-PCR |
| Ndufa5 | Ndufa5-p1 | CCTAGGCAATTCAGTATCCCTTACGC | KO isolation |
| | Ndufa5-p2 | CTCGAGGGTCTGGTTTCCAACAATAA | KO isolation |
| | Ndufa5-p3 | GTTAACGATGTTGACATGCACAGAAG | KO isolation |
| | Ndufa5-p4 | TTAATTAAGACAACTAGGAACCATCCAA | KO isolation |
| | Ndufa5-p5 | ATCAAACCCTGTACACCAAC | KO screening |
| | Ndufa5-p6 | TGCAACTCATCTGTCCAATA | KO screening |
| | Ndufa5-RTf | TGCTTCATCATGTTTTTGAG | RT-PCR |
| | Ndufa5-RTr | AGAGCCAGATCAACAAGAAA | RT-PCR |
| Hygromycin resistance cassette | HNZ Up rev | TGCGCAACTGTTGGGAAG | KO screening |
| | HN Down for | CCGCTGAAATCACCAGTCTC | KO screening |
| Actin | Actin2f | GCGAAGAGCGAGTATGACGAG | KO screening, RT-PCR |
| | Actin2r | AGCCACGAATCTAACTTGTGATG | KO screening, RT-PCR |
